# Cryo-EM structure of the receptor-activated TRPC5 ion channel at 2.9 angstrom resolution

**DOI:** 10.1101/467969

**Authors:** Jingjing Duan, Jian Li, Gui-Lan Chen, Bo Zeng, Kechen Xie, Xiaogang Peng, Wei Zhou, Jianing Zhong, Yixing Zhang, Jie Xu, Changhu Xue, Lan Zhu, Wei Liu, Xiao-Li Tian, Jianbin Wang, David E. Clapham, Zongli Li, Jin Zhang

**Affiliations:** School of Basic Medical Sciences, Nanchang University, Nanchang, Jiangxi, 330031, China.; Howard Hughes Medical Institute, Janelia Research Campus, Ashburn, VA 20147, USA; College of Pharmaceutical and Biological Engineering, Shenyang University of Chemical Technology, Shenyang 110142, China; Gannan Medical University, XueYuan Road 1#, Ganzhou, Jiangxi, 341000, China.; Key Laboratory of Medical Electrophysiology, Ministry of Education, and Institute of Cardiovascular Research, Southwest Medical University, Luzhou, Sichuan, 646000, China; The Key Laboratory of Molecular Medicine, the Second Affiliated Hospital of Nanchang University, Nanchang 330006, China.; College of Food Science and Engineering, Ocean University of China, Qingdao 266003, China.; School of Molecular Sciences and Biodesign Center for Applied Structural Discovery, Biodesign Institute, Arizona State University, Tempe, AZ 85287, USA.; Human Population Genetics, Human Aging Research Institute (HARI) and School of Life Science, Nanchang University, The Key Lab of Jiangxi Province for Human Aging, Nanchang, 330031, China.; Howard Hughes Medical Institute, Department of Biological Chemistry and Molecular Pharmacology, Harvard Medical School, Boston, MA 02115, USA.

**Author notes:** These authors contributed equally to this work (Jingjing Duan, Jian Li, Gui-Lan Chen).

## Abstract

The transient receptor potential canonical subfamily member 5 (TRPC5) is a non-selective calcium-permeant cation channel. As a depolarizing channel, its function is studied in the central nervous system and kidney. TRPC5 forms heteromultimers with TRPC1, but also forms homomultimers. It can be activated by reducing agents through reduction of the extracellular disulfide bond. Here we present the 2.9 Å resolution electron cryo-microscopy (cryo-EM) structure of TRPC5. The structure of TRPC5 in its apo state is partially open, which may be related to the weak activation of TRPC5 in response to extracellular pH. We also report the conserved negatively charged residues of the cation binding site located in the hydrophilic pocket between S2 and S3. Comparison of the TRPC5 structure to previously determined structures of other TRPC and TRP channels reveals differences in the extracellular pore domain and in the length of the S3 helix. Together, these results shed light on the structural features that contribute to the specific activation mechanism of the receptor-activated TRPC5.

## Introduction

The transient receptor potential canonical (TRPC) channels are Ca^2+^-permeant cation channels that are potentiated by stimulation of G protein-coupled receptors or receptor tyrosine kinases^1,2^. TRPC4 and TRPC5 share 65% sequence identity and exhibit similar functional properties when expressed as independent homomers. As homomultimers, the channel’s outward conductance is blocked in the 10-40 mV range^3,4^ by intracellular Mg^2+^ ^5^. TRPC5 is expressed in both excitable and non-excitable cells, primarily in the brain and kidney^6^. TRPC5 knock-out mice exhibit severe daily blood pressure fluctuations, and deficits in motor coordination and innate fear behavior^7-9^. Although contested^10^, TRPC5 inhibition may suppress the development of progressive kidney diseases^11^.

The cryo-EM revolution has enabled recent structural solutions for TRPC3, TRPC4, and TRPC6^12-14^. Here we present the structure of the mouse TRPC5 at pH 7.5 to an overall resolution of 2.9 Å. We contrast and compare the TRPC5 structure with other TRP channels to help understand the diverse functional and physiological roles of this ion channel family.

## Results

### Overall structure of the mouse TRPC5 tetrameric ion channel

A mouse TRPC5 construct lacking the C-terminal 210 residues (a.a. 1-765, excluding a.a. 766-975) yielded more protein after purification than that of the full-length construct. The baculovirus construct, consisting of a maltose binding protein (MBP) tag at the N terminus, was stably purified to homogeneity in *n*-dodecyl β-D-maltoside (DDM) and reconstituted in the amphipol poly (maleic anhydride-alt-1-decene) substituted with 3-(dimethylamino) propylamine (PMAL-C8; **Supplementary Fig. 1**). The amphipol reconstituted detergent-free protein was then negatively stained and analyzed by single-particle cryo-EM.

Single particle cryo-EM analyses of TRPC5 at an overall resolution of ∼ 2.9 Å was sufficient for *de novo* model building (**Supplementary Fig. 2**, **Supplementary Fig. 3, Supplementary Table 1**). Disordered regions led to poor densities for 7 residues in the S1-S2 loop, 28 residues in the distal N terminus, and 3 residues in the truncated distal C terminus. Similar to other solved TRP channel structures, TRPC5 forms a four-fold symmetric homotetramer (**Fig. 1a, b**) with dimensions of 100 Å × 100 Å × 120 Å. Each of the four monomers can be divided into a compact cytosolic domain and a transmembrane domain (TMD) (**Fig. 1)**. The cytosolic domain is composed of the N-terminal region with an ankyrin repeats domain (ARD) and seven α-helices (HLH); the C-terminal subdomain contains a connecting helix and a coiled-coil domain. The transmembrane domain (TMD) is composed of six α-helices (S1-S6), a TRP domain, and several small helices, including a pore helix, pre-S1 elbow, pre-S1 helix and an S2-S3 linker helix with connecting loops (**Fig. 1c, d)**.

**Figure 1.**
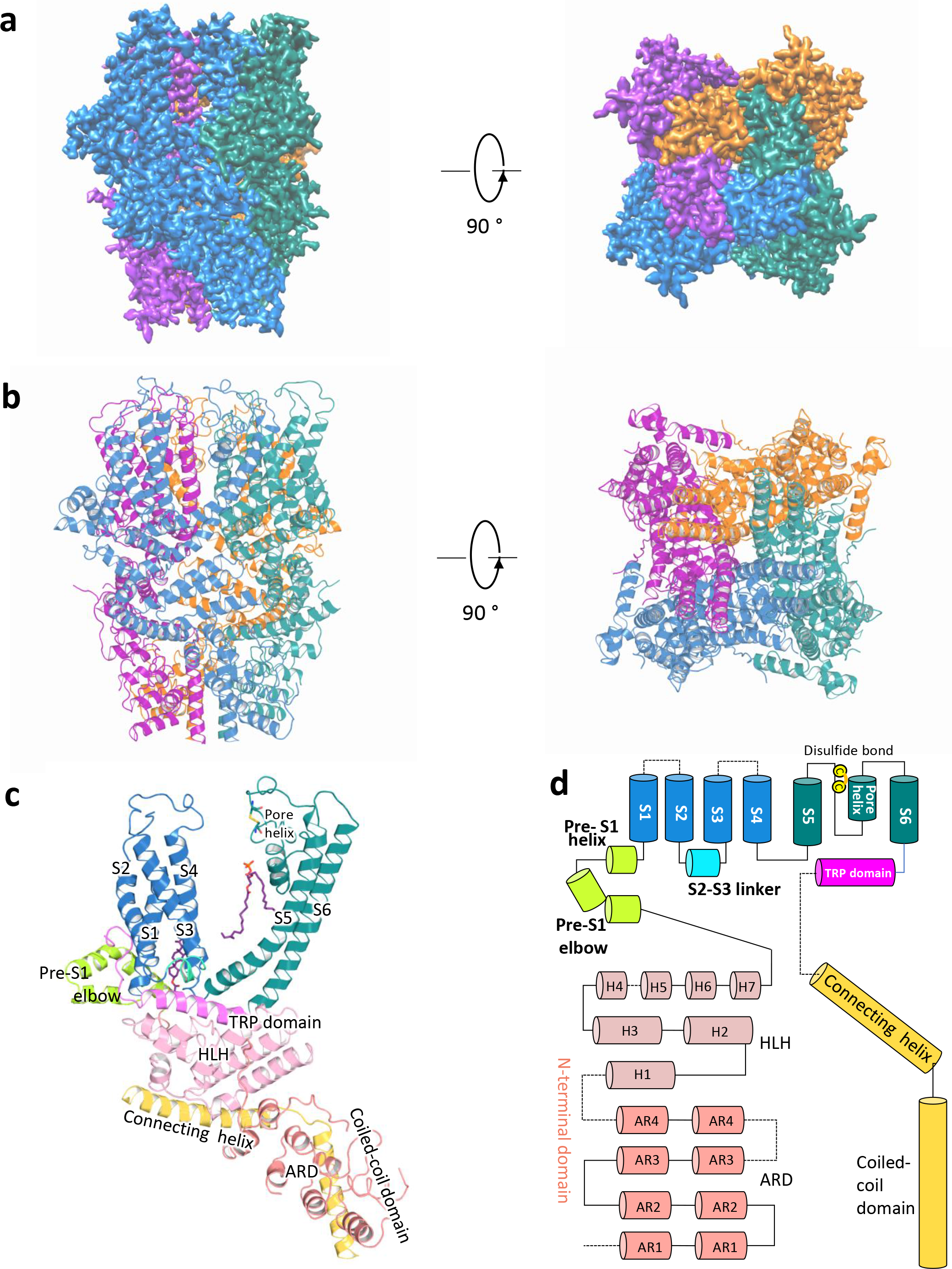
Overall structure of mouse TRPC5. **a**, Cryo-EM density map of mouse TRPC5 at 2.9 Å overall resolution with each monomer represented in different colors (left, side view and right, top view). **b**, Ribbon diagrams of the mouse TRPC5 model with the channel dimensions indicated. **c,** Ribbon diagrams depicting structural details of a single subunit. **d**, Linear diagram depicting the major structural domains of the TRPC5 monomer, color-coded to match the ribbon diagram in c.

Whole-cell patch-clamp recordings confirmed that the truncated TRPC5 construct used for cryo-EM analyses retained the primary electrophysiological properties of full-length TRPC5 (**Fig. 2**). Both the truncated and full-length constructs were activated by englerin A and inhibited by ML204, a TRPC4/5 blocker (**Fig. 2a, b**). No significant current was observed in cells transfected with empty vector (**Fig. 2c**).

**Figure 2.**
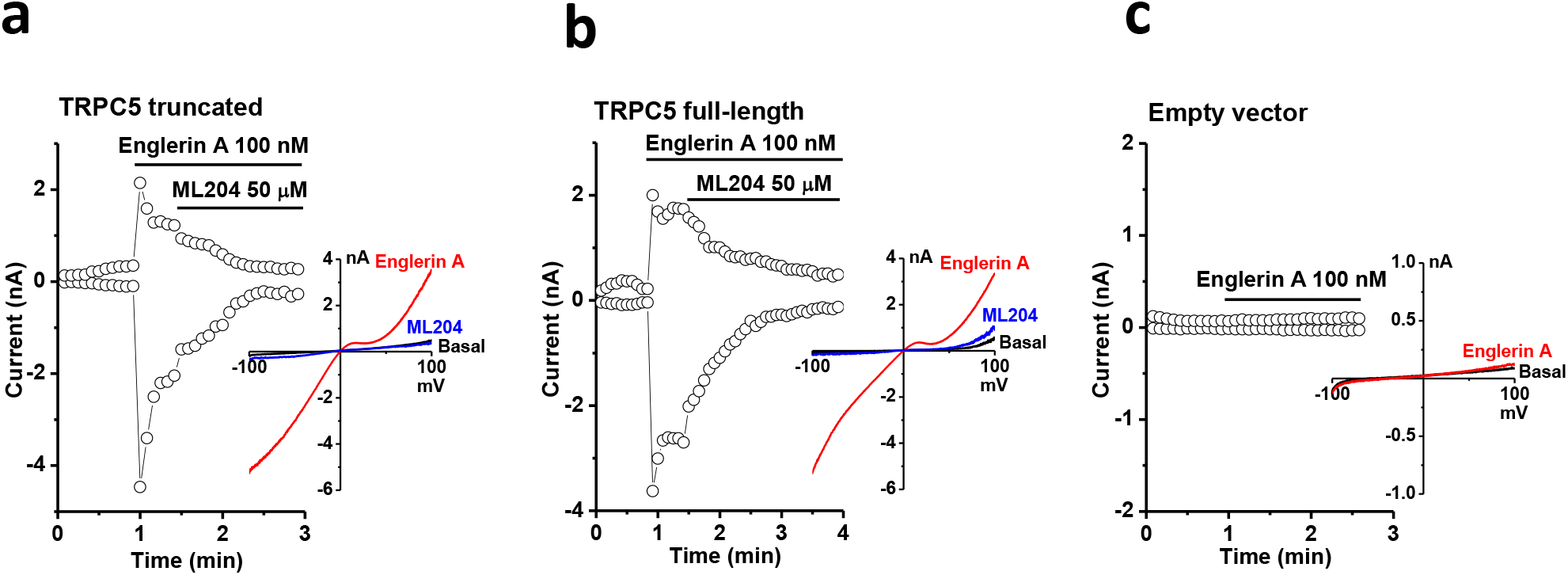
The truncated TRPC5 construct used for cryo-EM encodes a functional channel. Representative whole-cell patch clamp recordings and *I-V* relationships of **(a)** truncated TRPC5, **(b)** full-length TRPC5, and **(c)** empty vector-transfected HEK293 cells. The time course of currents measured at +80 and -80 mV and *I-V* relationships of the peak currents in different conditions are shown. Englerin A is an activator and ML204 is an inhibitor for TRPC4/5 channels.

### The ion conduction pore

TRPC4 and TRPC5 are the most closely-related TRPC proteins, with 65% amino acid identity at full length, and 100% identity in S5 (a.a. 502-542), the pore helix (a.a. 569-580), S6 (a.a. 596-651, including the lower gate) and the selectivity filter (a.a. 580-584) (**Supplementary Fig. 4**). The architecture of the ion conduction pathway is well conserved between these two channels (**Fig. 3a**). However, the calculated pore size of TRPC5 is wider than that of TRPC4 (**Fig. 3b)**, potentially due to the activation state of the purified protein. TRPC5’s motif ‘TRAIDEPNN’ (a.a. 544-552) forms a small helix in the middle of the extracellular loop connecting S5 and the pore helix (**Supplementary Fig. 5**). This unique helix, close to the disulfide bond and on the top of the pore helix (**Supplementary Fig. 5**), is distinct from the corresponding motif ‘ETKGLS’ of TRPC4. The TRPC5 motif contains 3 more amino acids, including one more negatively-charged residue, Asp548, and the hydrophobic residue, Ile547 (**Fig. 3c, red arrow**); it is thus potentially more attractive to cations entering the pore.

**Figure 3.**
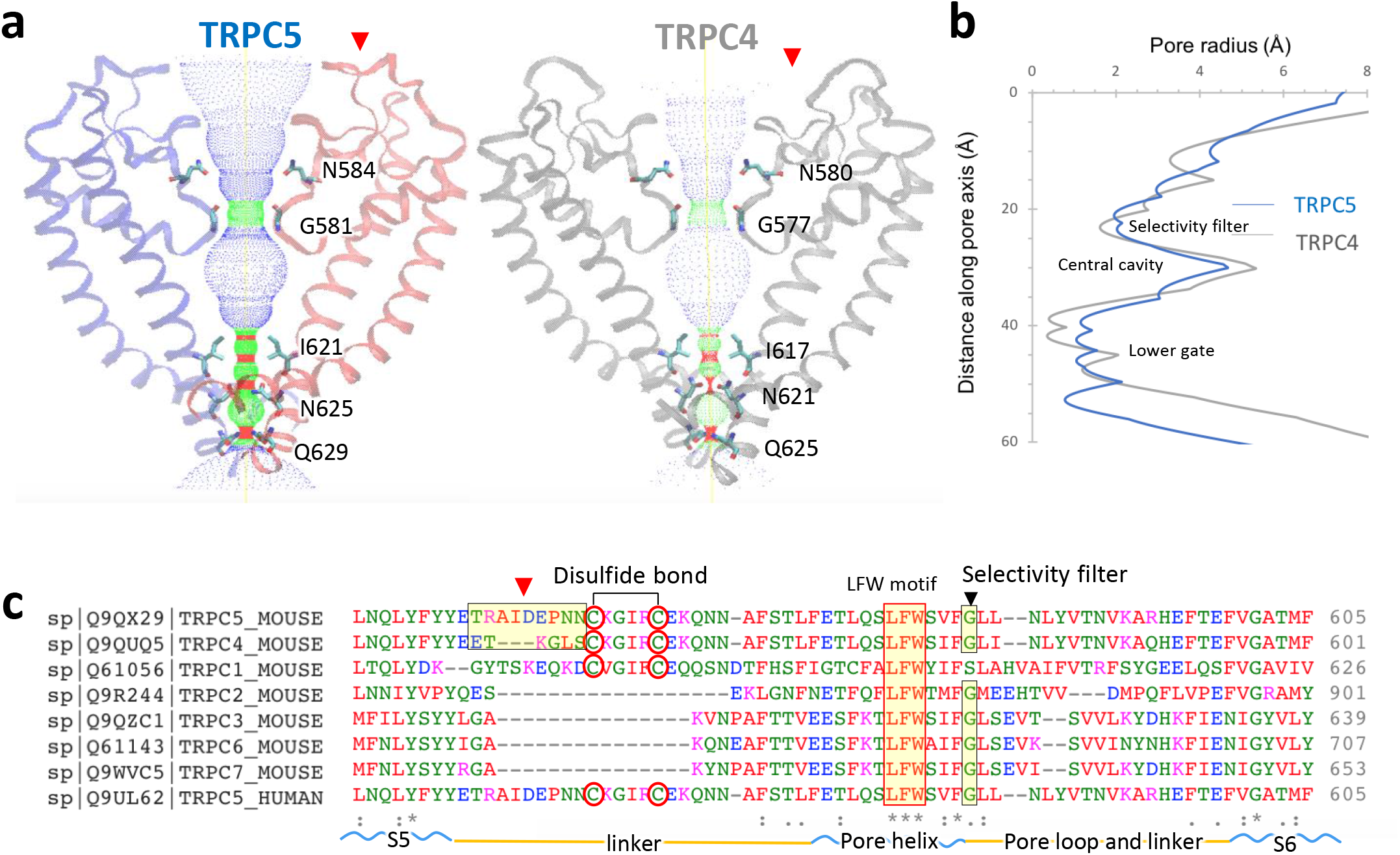
TRPC5 ion conduction pathway compared with TRPC4 and other TRPCs. **a**, Side view of TRPC5’s pore region with chains A and C compared with TRPC4 (gray). Ion conduction pathway shown as dots and mapped using HOLE with amino acid residues labeled. **b**, Pore radius along the central axis. The side chains of glycine form a narrow constriction at the selectivity filter. “INQ” motif forms the lower gate. **c**, Sequence of the mouse TRPC5 aligned to TRPC4 and other TRPC subfamily members (Clustal Omega) between S5 and S6 including the linker, pore helix, and pore loop. Regions corresponding to different extracellular pore domains are indicated by the red arrow. The two cysteines forming disulfide bonds, a conserved “LFW” motif, and the selectivity filter are highlighted.

Along with the pore, the filter/gate-forming residues of TRPC5 are identical to those of TRPC4. Gly581 marks a restriction point of 7.2 Å between diagonally opposed residues at the selectivity filter. “INQ” (Ile621, Asn625 and Gln629) defines a lower gate with a 4.9 Å constriction formed by the side chains of Gln629 (**Supplementary Fig. 6**); in TRPC4, the most restricted point is 3.6 Å between opposing Asn621 residues. Interestingly, the lower gate of TRPC5 appears to be partially open compared with the presumed closed state of TRPC4, despite similar isolation procedures and conditions. This may be related to the weak activation of TRPC5 compared to TRPC4 at pH 7.4^15^, suggesting a distinct activation mechanism for TRPC5 in response to extracellular pH.

### Structural comparisons between TRPC5 and other TRPC channels

Here we compare the architectural differences between the TRPC1/4/5 and TRPC3/6/7 channel subfamilies (**Fig. 4a**). An overlay of TRPC5 with the previously solved TRPC3, TRPC4, and TRPC6 channel structures shows relatively high spatial conservation of the 6-transmembrane bundle. Several intracellular features of TRPC channels are also preserved, including the ankyrin repeats, the pre-S1 elbow in the N-terminal domain, and the connecting helix running parallel to the membrane bilayer. A key feature of TRPC channels is the conserved LFW motif inside the pore domain (**Fig. 3c)**. In the TRPC5 structure, a π–π interaction between Phe576 and Trp577 stabilizes the key pore region, as is also seen in the available TRPC3/4/6 channel structures (**Fig. 4b**). Mutagenesis studies of the LFW motif in TRPC5 result in non-functional channels^16^.

**Figure 4.**
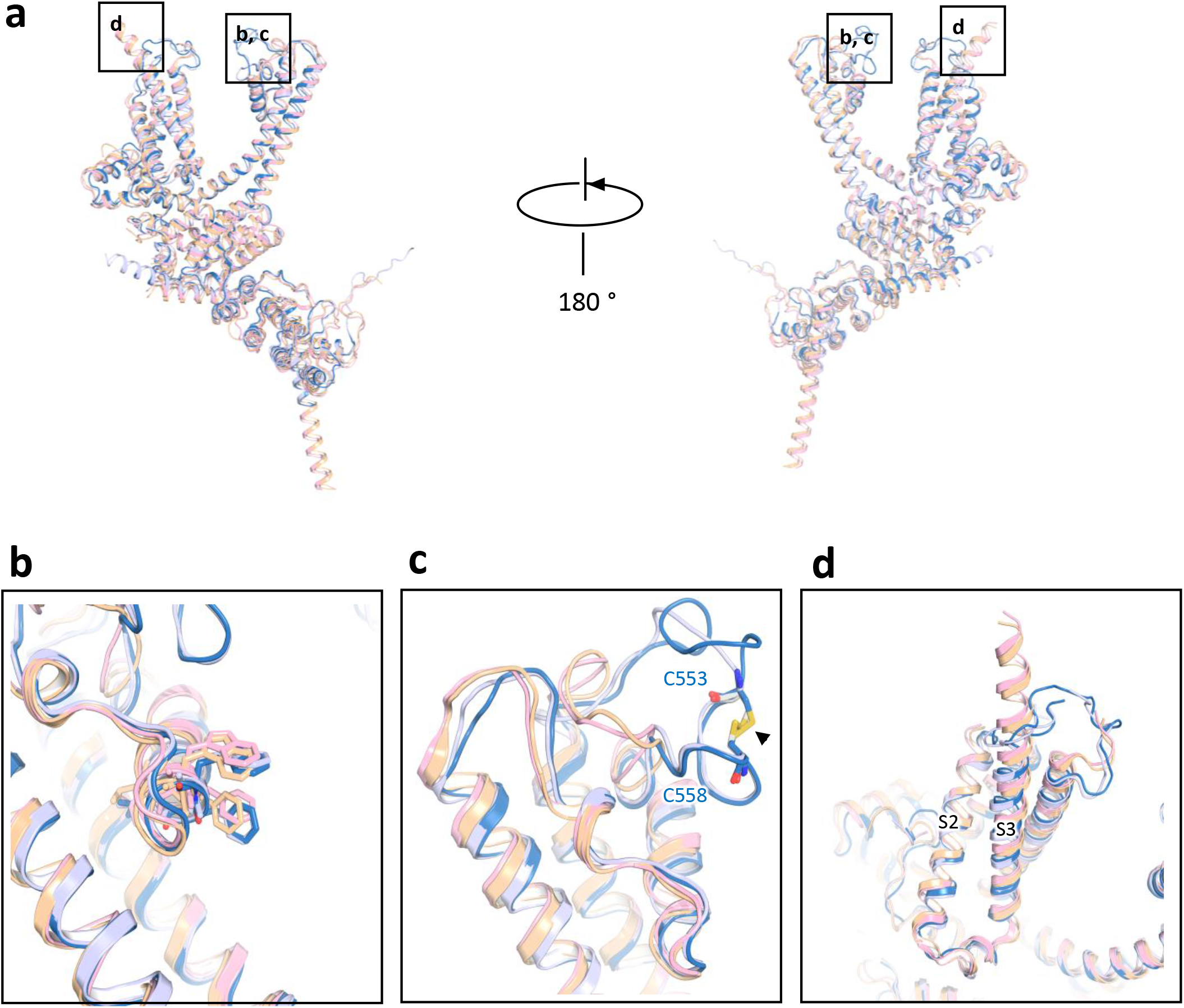
Comparison of the TRPC5 structure with other TRP channel structures. **a**, Superimposed side views of mouse TRPC5 subunit (blue) compared with other TRPC family members including mouse TRPC4 (PDB: 5Z96, gray)^37^, human TRPC3 (PDB: 5ZBG, pink)^12^, and human TRPC6 (PDB: 5YX9, yellow)^14^. **b**, The conserved “LFW” motif in the pore helix. **c**, Key pore loop disulfide bond between Cys553 and Cys548 in TRPC5 and the corresponding pore loop-disulfide bond in TRPC4. This disulfide bond is not present in TRPC3 or TRPC6 (black arrow); **d**, Differences in the organization of the S3 helix between the TRPC4/5 and TRPC3/6. The S3 helixes of TRPC3 and TRPC6 are longer than those of TRPC5 and TRPC4.

Despite the relatively high structural conservation, TRPC5’s transmembrane domain has several distinct features. Unlike TRPC3/6, TRPC4/5’s extracellular domains and long pore loops form intricate structures stabilized by a disulfide bond between Cys553 and Cys558. As the two cysteines in the pore region are well conserved in TRPC1/4/5 subfamily (**Fig. 3d**), the disulfide bond may be important to gating. Indeed, mutation of Cys553 or Cys558 to alanine activates TRPC5^17^ (**Fig. 4c**). Interestingly, although TRPC4 and TRPC5 have similar functional properties, the loop preceding TRPC5’s disulfide bond is distinct from that of TRPC4, with 3 extra residues (a.a. 546-548) in TRPC5 (**Fig. 3c**). Superimposition of the TRPC channels also reveals striking differences in the arrangement of S3 (**Fig. 4d**); the S3 helix is longer in TRPC3 and TRPC6 and protrudes into the extracellular space. In both TRPC4 and TRPC5 channels, the extracellular S3 region is shorter by four helical turns, limiting potential extracellular interactions.

### Cation- and lipid-binding sites

A strong non-protein peak is observed in the intracellular hydrophilic pocket between TRPC5’s S2 and S3 helices. Since Na^+^ was the only added cation in our purification buffer, this peak may represent a sodium ion, but we cannot rule out another contaminating cation, such as Ca^2+^. The cation is not in the putative ion conduction pathway; it is coordinated by hydrogen bonding with several highly conserved residues, including Glu418, Glu421, Asn436 and Asp439 (**Fig. 5a**). The geometry of the cation’s coordination in TRPC5 resembles that of the TRPC4 structure^18^. Structural alignment of these residues from TRPC family members reveals a well-preserved cation-binding site, including Ca^2+^ binding sites, in the TRPM4 and TRPM2 structures^19,20^. Small deviations are seen in TRPM4 and TRPC4; Glu421 in the S2 helix’s cytosolic end is substituted by a glutamine (**Fig. 5b, c**). A number of negatively charged residues, including Asp633, Asp636, Glu638, Asp652 and Glu653 in the TRP domain, as well as Asp424, Glu429 and Asp433 on S2-S3 linker, are aligned along a potential cation entry pathway (**Supplementary Fig. 7**). Asp652 and Glu653 in the TRP domain and negatively charged residues in the cytosolic region form a cation access pathway, which should facilitate cation entry to its binding site (**Supplementary Fig. 7c**). Further experiments are needed to determine if this represents a site for internal cationic potentiation or inhibition of TRPC5 activity.

**Figure 5.**
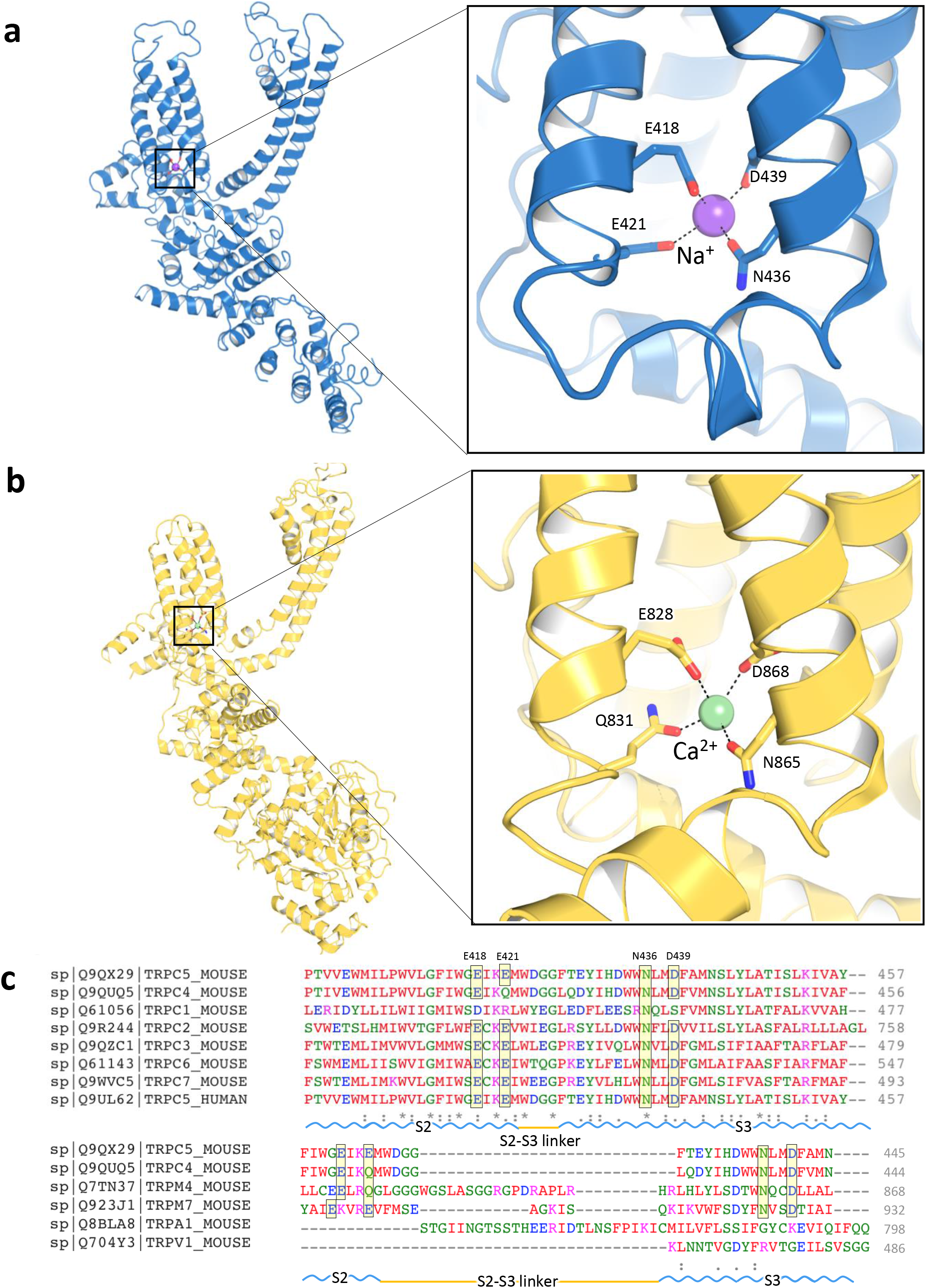
The cation binding site in TRPC5. **a**, A putative Na^+^ (purple sphere) on the cytosolic face is in the hydrophilic pocket of the S1-S4 domain, interacting with Glu418, Glu421, Asn436 and Asp439. **b**, Enlarged view of the putative Na^+^ binding site. Comparable Ca^2+^ (green sphere) binding site and an enlarged view of TRPM4 as “E/Q/N/D” (TRPC5: “E/E/N/D”). **c**, Sequence of the mTRPC5 aligned to mTRPC4 and other representative TRP members (Clustal Omega). The key residues are highlighted as “E/E/N/D”, which are conserved in TRPC2/3/5/6/7 and TRPM7 but as “E/Q/N/D” in TRPC4 and TRPM4.

Eight densities corresponding to lipid molecules were clearly resolved in the TRPC5 cryo-EM density and identified as four cholesteryl hemisuccinates (CHS, a cholesterol-mimicking artificial detergent molecule; purple) and four phospholipids (phosphatidic acid or ceramide-1-phosphate; yellow) (**Fig. 6a**). Each phospholipid is embedded in the gap between the monomeric subunits. The phospholipid interacts with the pore helix through its polar head and side chain oxygen with amino acids Phe576 and Trp577 (**Fig. 6b, c**). Four CHS heads face down at the interface with the plasma membrane, imbedding into the space among the S4, S5, S6 helices and the N-terminal domain (**Fig. 6b**). CHS stabilizes the domain interaction through interactions with Asn500 on the S4/S5 linker and with Trp315, Tyr316, and Trp322 in the TRP domain (**Fig. 6d**). The polar heads of the identified lipids interact with positively charged regions of the channel (**Supplementary Fig. 8**). Thus, we assume that *in vivo* phosphorylation or dephosphorylation, and the composition of membrane lipids, regulate ion conduction by altering channel topology.

**Figure 6.**
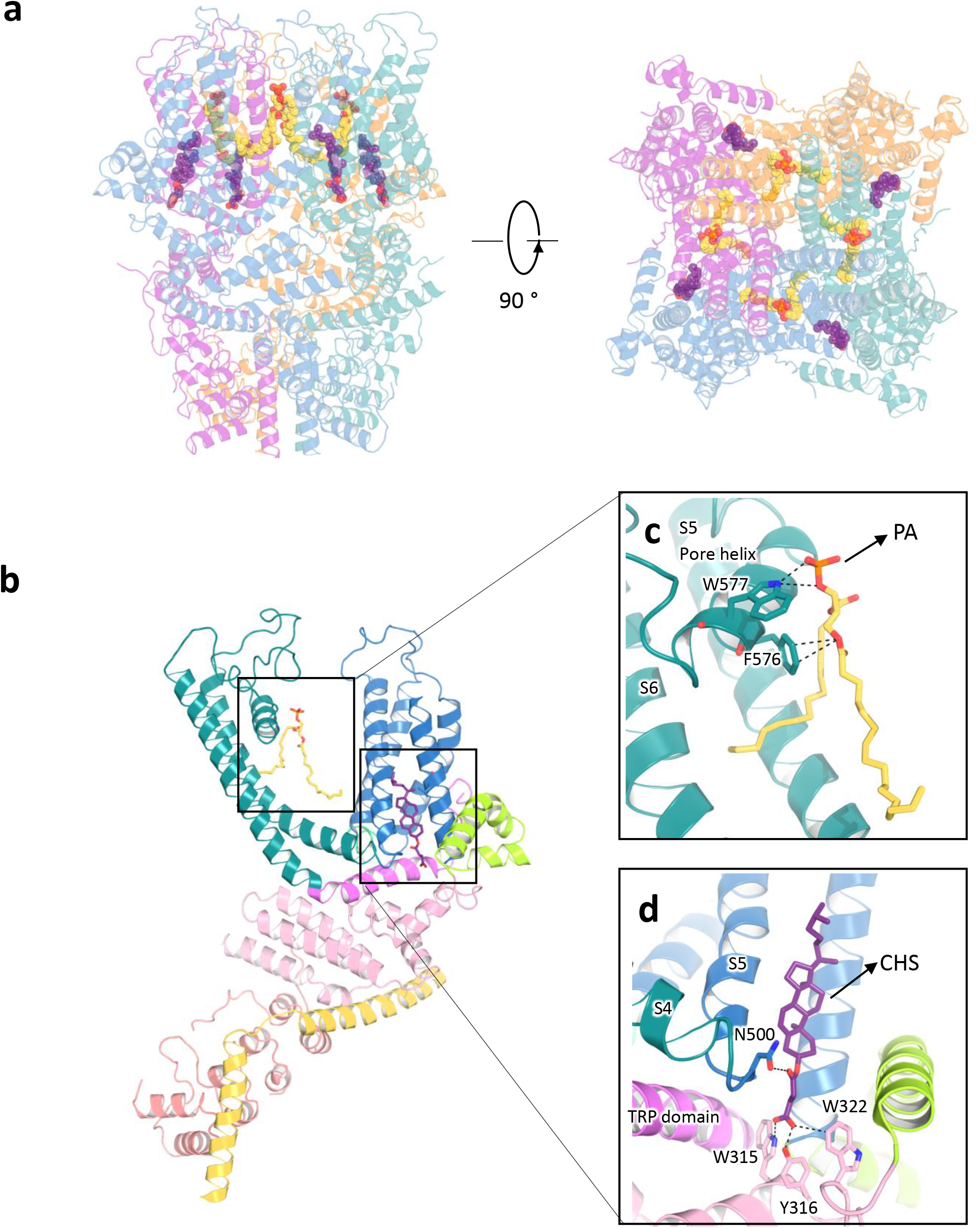
Lipid coordination in TRPC5. Lipid-channel interactions. **a**, Side and top views of ribbon diagrams of the TRPC5 tetramer: four cholesterol hemisuccinate (CHS) molecules and four phospholipids (potentially ceramide-1-phosphate, C1P, or phosphatidic acid, PA) are shown as spheres with purple or yellow carbons, respectively. **b**, Side views of each CHS and PA molecules per protomer. **c** and **d**, Ribbon diagram of the TRPC5 lipid-binding regions. **c**, PA interacts with the pore helix through Trp577 and Phe576. **d**, CHS, shown in purple, interacts with the S4/S5 linker at Asn500 and the N-terminal domains at Trp315, Tyr316, and Trp322.

## Discussion

Here we obtained and investigated the cryo-EM structure of TRPC5 at 2.9 Å resolution. The overall architecture of TRPC5 resembles that of other TRPC subfamily channels, but important differences in TM bundles and pore regions may help explain many of its unique functional features. Comparison of the structures of the open, or partially open, TRPC5 and closed TRPC3/6, suggest that the extracellular disulfide bond is a potential transducer of conformational changes that control the opening and closing of the pore domain. Another unique feature is the short S3 helix that dips into the hydrophobic region. Shift in this helical domain may be directly translated into the transmembrane domains.

One of the unique and interesting features of TPRC4/5 channels is that they are reversibly potentiated by micromolar concentrations of La^3+^ or Gd^3+^. Extensive mutagenesis studies suggest that the binding site is extracellular and between Glu543 and Glu595^21^. Mutation of glutamic acids Glu543 and Glu595/Glu598, in the extracellular mouth of the channel pore, to glutamine resulted in loss of potentiation. However, Glu543 at the end of the S5 helix, and Glu595 at the beginning of S6, are oriented such that cation binding is not possible. Since our structure was obtained in a divalent-free buffer, we can’t rule out the possibility that Ca^2+^ binding may lead to a conformational change in this region. (**Supplementary Fig. 9**).

Arg593, the arginine located in the pore loop near the proposed extracellular Gd^3+^ binding site, was reported to be critical for GPCR-G_q_-PLC-dependent gating of TRPC5^21^. An Arg593Ala mutation had reduced sensitivity to G_q_-PLC activation, which can be explained by our structure. Arg593 interacts via a salt bridge with Glu598 and forms a hydrogen bond with the Val590 side chain. Breaking this electrostatic interaction may lead to unfolding of this pore region. In addition, Arg593 is the only positively charged residue in the corresponding positions in TRPC channels (glycine in TRPC1, glutamine in TRPC4, asparagine in TRPC6, or negatively-charged aspartic acid in TRPC7). One possibility that Arg593 may serve as a molecular fulcrum, allowing the efficient transmission of gating force to TRPC5’s pore helix-loop.

Another unique feature of TPRC4/5 channels is their unusual current-voltage (*I-V*) relationship, in particular the flat segment in the 10-40 mV range attributed to Mg^2+^ block of outward current^2^. TRPC5’s S6 terminal Asp633 residue is close to the intracellular mouth of channel pore and was proposed as the site for block of the ion conduction pathway upon binding Mg^2+^, whereas the downstream Asp636 did not affect Mg^2+^ block^22^. In our structure, Asp633 is situated much closer to the channel pore than Asp636 (**Supplementary Fig. 10**), which supports it being a Mg^2+^ block site. However, because we used a divalent-free buffer for cryo-EM, direct evidence for Mg^2+^-binding on Asp633 and restriction of the channel pore should be studied in proteins purified in Mg^2+^-containing solution.

In TRPV channels, as in many other 6TM channels, the S4-S5 linker is crucial to channel gating^23-25^. Consistent with this, a single point mutation in TRPC4/5’s S4-S5 linker, Gly504Ser, fully activates it. Introduction of a second mutation (Ser623Ala) into TRPC4 Gly503Ser suppressed the constitutive activation and partially rescued its function^26^. These results indicate that the S4–S5 linker is a critical constituent of TRPC4/C5 channel gating and that disruption disinhibits gating control by the sensor domain.

Both the TRPC5 structure in the present study and our recently published TRPC4 structure^18^ were obtained from protein samples prepared in the same divalent-free solution at pH 7.5. TRPC5’s wider pore could reflect that an open state is preferred under these conditions. In our patch-clamp experiments, TRPC5 always exhibited small but consistent basal activity, while TRPC4 was inactive without drug or receptor stimulation. This agrees with a previous study that TRPC5 is partially open at pH 7.4, while TRPC4 is closed^15^. The wider pore size of TRPC5 may allow small number of cations to pass through under resting conditions and suggest that the current TRPC5 structure is a partially open state of the channel.

In summary, our structure enables new comparisons between the different subgroups of the available TRPC channels. Our findings provide structural insights into conservation and divergence of TRPC channels, but do not explain the mechanisms regulating GPCR-G_q_-PLC activation of a receptor-operated channel. Finally, the findings have broad implications for understanding the structural basis underlying epitope antibody selectivity, and can facilitate rational, structure-based, approaches for the design of subtype-selective TRPC ligands as new therapeutic agents.

## Materials and Methods

### Protein expression and purification

A synthetic gene fragment encoding residues 1 to 765 (excluding a.a. 766-975) of mouse TRPC5 was cloned into the pEGBacMam vector. The resulting protein contains a maltose binding protein (MBP) tag on its N terminus. P3 baculoviruses were produced in the Bac-to-Bac Baculovirus Expression System (Invitrogen). HEK293S GnTI^-^ (from ATCC) cells were infected with 10% (v/v) P3 baculovirus at a density of 2.0 - 3.0 × 10^6^ cells/ml for protein expression at 37°C. After 24h, 10 mM sodium butyrate was added, and the temperature reduced to 30°C for 72h before harvesting. Cells were gently disrupted and resuspended in a solution containing 30 mM HEPES, 150 mM NaCl, 1 mM dithiothreitol (DTT), pH 7.5 with EDTA-free protease inhibitor cocktail (Roche). Cells were solubilized for 2-3h in a solution containing 1.0% (w/v) N-dodecyl-beta-D-maltopyranoside (DDM, Affymetrix), 0.1% (w/v) cholesteryl hemisuccinate (CHS, Sigma), 30 mM HEPES, 150 mM NaCl, 1 mM DTT, pH 7.5 with EDTA-free protease inhibitor cocktail (Roche). After 30 min, the cell lysate was then centrifuged for 60 min at 100,000×*g* and the supernatant incubated in amylose resin (New England BioLabs) at 4°C overnight. The resin was washed with 20 column volumes of wash buffer [25 mM HEPES, 150 mM NaCl, 0.1% (w/v) digitonin, 0.01% (w/v) CHS, 1 mM DTT; pH 7.5 with EDTA-free protease inhibitor cocktail (Roche)]. The protein was eluted with 4 column volumes of wash buffer with 40 mM maltose. The protein was then concentrated to 0.5 ml and mixed with PMAL-C8 (Anatrace) at 1:4 (w/w) with gentle agitation for 4h. Bio-Beads SM-2 (50 mg/ml reconstitution mixture, Bio-Rad) were added to remove additional detergent; a disposable poly-prep column was used to remove Bio-Beads. After incubation at 4°C overnight, the protein was further purified on a Superose 6 size exclusion column in 25 mM HEPES, 150 mM NaCl, 1 mM DTT; pH 7.5. All purification procedures were carried out either on ice or at 4 °C. The peak fractions corresponding to tetrameric TRPC5 was concentrated to 7.0 mg/ml and used for preparation of cryo-EM sample grids.

### Electron microscopy data collection

3.5µl of purified TRPC5 protein sample in PMAL-C8 at 7.0 mg/ml was applied to glow-discharged Quantifoil R1.2/1.3 holey carbon 400 mesh copper grids (Quantifoil). Grids were blotted for 7s at 100% humidity and flash frozen by liquid nitrogen-cooled liquid ethane using a FEI Vitrobot Mark I (FEI). The grid was then loaded onto FEI TF30 Polara electron microscope operated at 300 kV accelerating voltage. Image stacks were recorded on a Gatan K2 Summit direct detector (Gatan) set in super-resolution counting mode using SerialEM^27^, with a defocus range between 1.5 and 3.0 μm. The electron dose was set to 8 e-/physical pixel/s and the sub-frame time to 0.2 s. A total exposure time of 8s resulted in 40 sub-frames per image stack. The total electron dose was 42.3 e-/Å^2^ (∼1.1 e-/Å2 per sub-frame).

### Image processing and 3D reconstruction

Image stacks were gain-normalized and binned by 2x to a pixel size of 1.23Å prior to drift and local movement correction using motionCor2^28^, resulted in the sums of all frames of each image stack with and without dose-weighting. The dose-weighting sum were binned by 8x and subject to visual inspection, images of poor quality were removed before particle picking. Particle picking and subsequent bad particle elimination through 2D classification was performed using Python scripts/programs^29^ with minor modifications in the 8x binned images. The selected 2D class averages were used to build an initial model using the common lines approach implemented in SPIDER^30^ through Maofu Liao’s Python scripts^29^, which was applied to later 3D classification using RELION^31^. Contrast transfer function (CTF) parameters were estimated using CTFFIND4^32^ using the sum of all frames without dose-weighting. Quality particle images were then boxed out from the dose-weighted sum of all 40 frames and subjected to RELION 3D classification. RELION 3D refinements were then performed on selected classes for the final map. The resolution of this map was further improved by using the sum of sub-frames 1-14. The number of particles in each dataset and other details related to data processing are summarized in Table 1.

### Model building, refinement, and validation

At 2.9 Å resolution, the cryo-EM map was of sufficient quality for de novo atomic model building in Coot^33^. Amino acid assignment was achieved based mainly on the clearly defined densities for bulky residues (Phe, Trp, Tyr, and Arg) and the absence of side chain densities for glycine residues. The atomic model was further visualized in COOT; a few residues with side chains moving out of the density during the refinement were fixed manually, followed by further refinement. The TRPC5 model was then subjected to global refinement and minimization in real space using the module ‘phenix.real_space_refine’ in PHENIX^34^. The geometries of the model were assessed using Mo Probity^35^ in the ‘comprehensive model validation’ section of PHENIX. The final model exhibited good geometry as indicated by the Ramachandran plot (preferred region, 98.25%; allowed region, 1.75%; outliers, 0%). The pore radius was calculated using HOLE^36^.

## Electrophysiology

The full-length, truncated TRPC5 constructs or empty vector were transfected into HEK 293T (from ATCC) cells together with an mCherry plasmid. Cells with red fluorescence were selected for whole-cell patch recordings (HEKA EPC10 USB amplifier, Patchmaster 2.90 software). A 1-s ramp protocol from –100 mV to +100 mV was applied at a frequency of 0.2 Hz. Signals were sampled at 10 kHz and filtered at 3 kHz. The pipette solution contained (in mM): 140 CsCl, 1 MgCl_2_, 0.03 CaCl_2_, 0.05 EGTA, 10 HEPES, and the pH was titrated to 7.2 using CsOH. The standard bath solution contained (in mM): 140 NaCl, 5 KCl, 1 MgCl_2_, 2 CaCl_2_, 10 HEPES, 10 D-Glucose, and the pH was adjusted to 7.4 with NaOH. The recording chamber (150 µl) was perfused at a rate of ∼2 ml/min. All recordings were performed at room temperature.

## Acknowledgements

We thank Dr. Steve Harrison and the Cryo-EM Facility (Harvard Medical School) for use of their microscopes. We thank Dr. Maofu Liao for providing the Python scripts and help in image processing. J.Z. was supported by the Thousand Young Talents Program of China, National Natural Science Foundation of China (Grant No. 31770795) and Jiangxi Province Natural Science Foundation (Grant No. 20181ACB20014). J.L. was supported by the National Natural Science Foundation of China (Grant No. 81402850). Functional studies in this project were supported by the National Natural Science Foundation of China (31300949 to B.Z. and 31300965 to G.L.C.)

## Author Contributions

J.Z., J.D. and D.E.C. designed the project. J.Z. and J.D. designed and made constructs for BacMam expression and determined the conditions used to enhance protein stability. J.Z. purified the protein. Z.L. carried out detailed cryo-EM experiments, including data acquisition and processing. J.L. and J.Z. built the atomic model on the basis of cryo-EM maps. G.L.C., B.Z. and K.C. performed functional studies. X. P., J. Z., Y.Z., X. J., C.X., L.Z., W.L., X.L.T. and J.W. assisted with protein purification and the mutation of TRPC5 constructs for functional studies. J.D. and J.Z. drafted the initial manuscript. All authors contributed to structure analysis/interpretation and manuscript revision. J.Z., D.E.C. and Z.L. initiated the project, planned and analyzed experiments and supervised the research.

The authors declare no competing financial interests.

Data deposition: Cryo-EM electron density map of the mouse TRPC5 has been deposited in the Electron Microscopy Data Bank, https://www.ebi.ac.uk/pdbe/emdb/ (accession number EMD-9615), and the fitted coordinate has been deposited in the Protein Data Bank, www.pdb.org (PDB ID code 6AEI).

## Supplementary Figures

**Supplementary Figure 1. Biochemical characterization of the TRPC5 construct.**

Size exclusion chromatography of TRPC5 proteins. Void volume (V_0_) and the peaks corresponding to tetrameric TRPC5 and PMAL-C8 are indicated. Protein samples of the indicated TRPC5 protein fraction were subjected to SDS-PAGE and Coomassie-blue staining.

**Supplementary Figure 2. Flow chart for TRPC5 cryo-EM data processing.**

**a**, Representative image of the purified TRPC5 protein, 2D class averages of TRPC5 particles, side views of the 3D reconstructions from RELION 3D classification, and final 3D reconstructions from 3D auto-refinement. **b**, Fourier shell correlation (FSC) curve for the 3D reconstruction (marked at overall 2.9 Å resolution). **c**, Local resolution estimation from ResMap^38^ and, **d**, Euler distribution plot of particles used in the final three-dimensional reconstruction. The length of the rod is proportional to the number of particles in that view, with regions in red denoting the views containing the highest number of particles.

**Supplementary Figure 3. Cryo-EM densities of selected regions of TRPC5.**

Density map showing the transmembrane helices (S1-S6), an ankyrin repeat (AR), N-terminal helix, TRP domain, pore helix, connecting helix, coiled-coil helix, and lipids. The maps were contoured at a level of 3.0 σ, carve = 2. Ankyrin repeats, AR: T144-K164, N helix: P216-E234, S1: N355-Q386, S2: V402-W423, S3: F427-Y457, S4: F474-F500, S5: H502-Y542, pore helix: F569-F580, S6: F596-F651, connecting helix: L707-N735 and coiled-coil domain: E740-G761.

**Supplementary Figure 4. Sequence alignment of TRPC subfamily members.**

Sequence of the full-length mouse TRPC5 aligned to other TRPC subfamily members (Clustal Omega); key residues indicated. Regions corresponding to the putative Na^+^ binding sites are labeled. The selectivity filter, lower gate, and two cysteines forming disulfide bonds are highlighted.

**Supplementary Figure 5. Pore loop structures of known TRP channels**

**a-h**, comparison of the pore loop in TRPC5 (**a**, PDB accession number 6AEI) with other known TRP channel, including TRPC4 (**b**, PDB accession number 5Z96), TRPC3 (**c**, PDB accession number 5ZBG), TRPC6 (**d**, PDB accession number 5YX9), TRPM4 (**e**, PDB accession number 6BWI), TRPM7 (**f**, PDB accession number 6BWD), TRPA1 (**g**, PDB accession number 3J9P) and TRPV1 (**h**, PDB accession number 5IRX). i, Sequence alignment of the pore loop in known TRP channels.

**Supplementary Figure 6. Comparison of ion conducting pathways between TRPC5 and TRPC4.**

Comparison of ion conduction pathways of **a**, TRPC5, and **b**, TRPC4 (PDB: 5Z96). Distances between specific side chains along the pore and the key residues are labeled.

**Supplementary Figure 7. The predicted Na^+^ binding sites.**

Side and top views of electrostatic maps of predicted Na^+^ binding pockets in TRPC4; **a,** Tetramer and **b,** Monomer. Gray dots highlight possible pathways for Na^+^ entry. The surface is colored according to the calculated electrostatic potential, revealing the tetrameric distribution of charge. Blue indicates positive potential, red negative potential, and transparent white, neutral potential. **c,** A number of negatively charged residues, including Asp633, Asp636, Glu638, Asp652 and Glu653 on the TRP domain, as well as Asp424, Glu429 and Asp433 on S2-S3 linker, are aligned along the cation entry pathway. **d,** The bottom view of the cytoplasmic cavity for cation.

**Supplementary Figure 8. Lipid-channel interactions.**

**a**, Side view of TRPC5 monomer. CHS (purple) and PA (yellow) molecules are shown as sticks. **b**, Electrostatic maps of the lipids binding sites.

**Supplementary Figure 9. Poor loop interactions**

Key residues located in the pore loop are shown in stick representation. Arg593 forms a hydrogen bond with Glu598.

**Supplementary Figure 10. Intracellular mouth of the channel pore**

Asp633 and Asp636 residues are shown in stick representation. Na^+^ bound between S2 and S3 are shown as purple spheres. Carbon and oxygen atoms of residues are shown in green and red, respectively.

